# SOAPTyping: an open-source and cross-platform tool for Sanger sequence-based typing for HLA class I and II alleles

**DOI:** 10.1101/674648

**Authors:** Yong Zhang, Yongsheng Chen, Huixin Xu, Junbin Fang, Zijian Zhao, Weipeng Hu, Xiaoqin Yang, Jia Ye, Yun Cheng, Jiayin Wang, Jian Wang, Huanming Yang, Jing Yan, Lin Fang

## Abstract

**Summary:** The human leukocyte antigen (HLA) gene family plays a key role in the immune response and thus is crucial in many biomedical and clinical settings. Utilizing Sanger sequencing - the gold standard technology for HLA typing – enables accurate identification of HLA alleles with high-resolution. However, there exists a current hurdle that only commercial software such as UType, SBT-Assign and SBTEngine, instead of any open source tools could be applied to perform HLA typing based on Sanger sequencing. To fill the gap, we developed a stand-alone, open-source and cross-platform software, known as SOAPTyping, for Sanger-based typing in HLA class I and II alleles.

**Availability and implementation:** SOAPTyping is implemented in C++ language and Qt framework, which is supported on Windows, Mac and Linux. Source code and detailed documentation are accessible via the project GitHub page: https://github.com/BGI-flexlab/SOAPTyping.

**Supplementary information:** Supplementary data are available at Bioinformatics online.

## INTRODUCTION

Human leukocyte antigens (HLA), encoded on 6p21.3, make up the human major histocompatibility complex (MHC) regions with high polymorphism and are featured in the immunity system (Dendrou et al., 2018). Accurate HLA allele determination (‘HLA Typing’) is potent and crucial in various biomedical and clinical processes, especially in the field of solid organ and bone marrow transplantation (Mahdi, 2013). Sequence Based Typing (SBT), including Sanger sequence-based typing (SSBT) and next-generation sequence (NGS) typing, is widely used for high-resolution four-digit allele level identification of HLA class I and II alleles (Erlich, 2012). Advantaging in producing the sequenced DNA in contiguous form, SSBT serves as the gold standard for HLA typing, which applies polymerase chain reaction (PCR) to amplify loci of targets while utilizing Sanger sequencing and related software to determine the nucleotide sequence of the PCR product. Sanger sequencing sometimes rises phase ambiguities due to multiple polymorphisms shared between alleles, which requires further steps using group specific sequencing primers (GSSP).

While SSBT is reliable and routine for clinical use, there are no open-source tools currently available but only commercial and Windows-supported software, such as UType (Life Technologies. Brown Deer, WI), SBT-Assign (Conexio, San Francisco, CA) and SBTEngine (GenDx, Utrecht, Netherlands), to perform sequence analysis and allele assignments for SSBT, and thus limits its application. Hence, SOAPTyping was developed as a fast, accurate and effective cross-platform software with user-friendly interface for SSBT in HLA class I and II alleles. Supported on Windows, Mac and Linux, SOAPTyping also provides a neat and interactive user interface and generates specialized report format. No proficient computer skills are required for users to effectively complete the analysis with a comprehensible protocol and produce accurate results. SOAPTyping also integrates group specific sequencing primers (GSSP) prediction system to resolve the alleles ambiguity. SOAPTyping is open source and freely available at https://github.com/BGI-flexlab/SOAPTyping.

## IMPLEMENTATIONS

SOAPTyping is a flexible and powerful application implemented in C++ with its user-friendly interface developed in Qt framework, which is supported on Windows, Mac and Linux. SOAPTyping is capable of analyzing loci located in HLA class I (A, B, C and G) and II (DR-, DQ- and DP-) genes (Table S1). It mainly comprises modules specialized for database, backend analysis and visualization.

### Database

SOAPTyping offers database update functions to cater to the frequently updated HLA alleles. Nucleotide sequence alignments files of the IMGT/HLA database (Robinson et al., 2015) were applied to perform sequence format conversion with the scripts provided on our website, such files ending up with storage in the static database to serve as the reference of alignments. GSSP prediction system is available to resolve the ambiguity caused by phase problems, that GSSP binds to only one of the two alleles present in the DNA sample aiding the determination of the final HLA typing. Involved database could be manually prepared for updates by following instructions in the supplementary materials (supplementary Section 2.9).

### Backend analysis

Sequences derived from the input ABIF files were called homozygotes or heterozygotes. After being presented as lists of degenerate bases, sequences are aligned to the consensus sequences and alleles in the IMGT/HLA database to assign the eligible allele pairs with utilization of a modified semi-global alignment method (supplementary Section 1.3). SOAPTyping produces a standardized output following nomenclature of HLA alleles (Marsh et al., 2010).

### Visualization

As shown in Figure 1, the results are presented in a main window of SOAPTyping consists of panes of Toolbar, Base Navigator, Sequence Display, Sample List, Allele Match List and Electropherogram Display.

**Figure 1.**
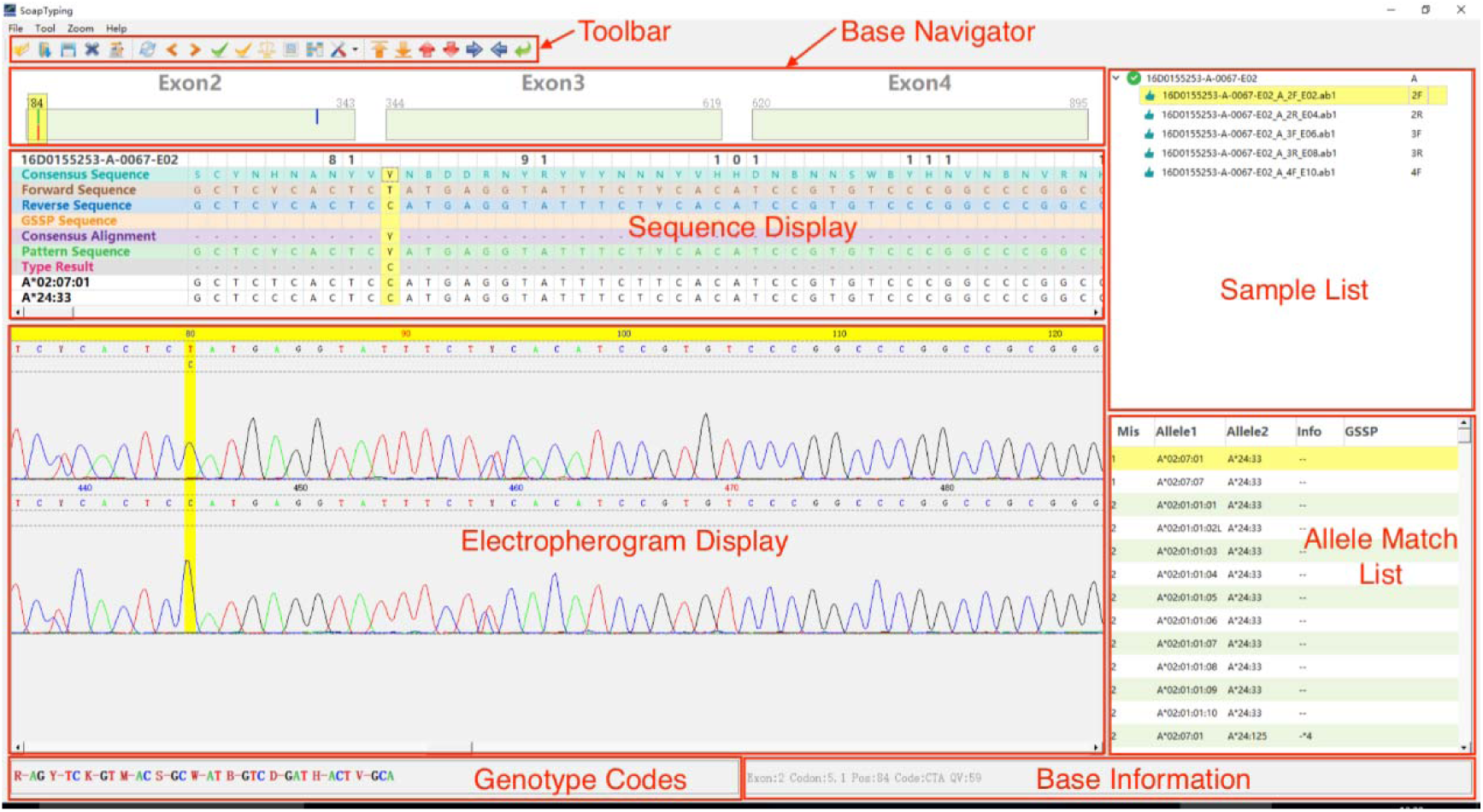
Main window of SOAPTyping. The pane of Sample List displays input files as a tree structure based on samples’ name. The pane of Base Navigator highlights mismatched positions so that users can skip to such positions quickly by clicking on the color bar. The pane of Allele Match List displays possible typing results sorted following the order of number of mismatched sites. The pane of Sequence Display, from top to bottom, is comprised of server tracks including ‘Sample and Position’, ‘Consensus Sequence’, ‘Forward Sequence’, ‘Reverse Sequence’, ‘GSSP Sequence’, ‘Consensus Alignment’, ‘Pattern Sequence’, ‘Type Result’ and sequences of the allele pair. The pane of Electropherogram Display displays the electropherogram of the forward sequence, the reverse sequence and the GSSP sequence, so that users can edit bases in this region. The pane of Toolbar integrates some useful functions and information, such as import and export reports.

### Best practices / proposed workflow

SOAPTyping works on chromatogram files with the format of ABIF, including .ab1 and .fsa files, which are generated from Sanger sequencing by ABI Genetic Analyzer Software (Applied Biosystems, Foster City, CA). Top candidate allele pair matches are presented in the Allele Match List. If necessary, users could manually review and edit marked positions which result from discrepant sites between forward and reverse sequences or mismatches with consensus sequence(s) till completion of at least one trace with ‘0’ mismatch in the Allele Match List. Best practices and proposed workflow are provided in Figure S3 and supplementary Section 2 to facilitate and guide efficient use of SOAPTyping.

## RESULTS

To verify the accuracy of SOAPTyping, our test data contains 36 samples initiated for external quality assessments with the University of California Los Angeles (UCLA) International HLA DNA Exchange (Los Angeles, CA, USA). Genomic DNAs with known HLA typing results were obtained from UCLA and amplified using locus-specific primers. The PCR products were directly sequenced in HLA-A, -B, -C, -DRB1 and -DQB1 (Table S1) using a 3730XL DNA Analyzer (Applied Biosystems, Foster City, CA). Sequencing reaction was performed using the BigDye® Terminator v3.1 Cycle Sequencing Ready Reaction Kit (Applied Biosystems). The sequence was analyzed with SOAPTyping and the typing results were compared to the consensus based on high resolution provided by UCLA. The consistency of SOAPTyping in typing HLA alleles at four-digit was verified to be accurate at the level of 100% (36/36) for HLA-A, 100% (36/36) for HLA-B, 100% (36/36) for HLA-C, and 100% (36/36) for HLA-DRB1, 100% (36/36) for HLA-DQB1. The detailed results of 36 tested samples were shown in Table S13. The test data have been deposited in the CNSA (https://db.cngb.org/cnsa/) of CNGBdb with an accession code CNP0000512.

## DISCUSSIONS

SOAPTyping was introduced in this article as the first open-source and cross-platform HLA typing software with the capability of producing high-resolution HLA typing predictions from Sanger sequence data. While high consistency with other commercial typing software is achieved comparing to actual HLA typing results, we demonstrated that SOAPTyping could be efficiently and effectively applied to practical use while some augmentation will still be anticipated in the future. With the challenge of upscaling of the HLA alleles in the IMGT/HLA database, future improvements of the efficiency of searching for the candidate allele pairs are needed to enhance its performance. Optimum search strategies will be required to develop while maintaining accuracy of typing results with at least four-digit resolution.

## Acknowledgements

We would also like to thank Taipu Lin, Jason Chen and Yuchong Tan for useful seggustions on the writing of the paper.

