## Supplemental Materials for "SOAPTyping: an open-source and cross-platform tool for Sanger sequence-based typing for HLA class I and II alleles"

SOAPTyping Supplement materials

### 1. Software Architecture

SOAPTyping is capable of analyzing loci located in HLA class I (A, B, C and G) and II (DRB1, DRB3, DRB4, DRB5, DPA1, DQA1, DQB1 and DPB1) genes (Table S1). SOAPTyping includes three modules as illustrated in the Figure S1, which are modules for (i) visualization, (ii) backend analysis, and (iii) database. These modules are described in further details in the following part.

Table S1. HLA molecules and the respective exon regions that can be analyzed by SOAPTyping

| Genes | Exons | Exons in the test data |
| --- | --- | --- |
| HLA-A | 1,2,3,4,5,6 | 2,3,4 |
| HLA-B | 1,2,3,4,5 | 2,3,4 |
| HLA-C | 1,2,3,4,5,6,7 | 2,3,4 |
| HLA-DRB1 | 1,2,3,4 | 2,3 |
| HLA-DRB3,4,5 | 2,3 | NA |
| HLA-DQA1 | 1,2,3,4, | NA |
| HLA-DQB1 | 1,2,3,4 | 2,3 |
| HLA-DPB1 | 1,2,3,4 | NA |
| HLA-G | 2,3,4 | NA |
| HLA-DPA1 | 1,2,3,4 | NA |

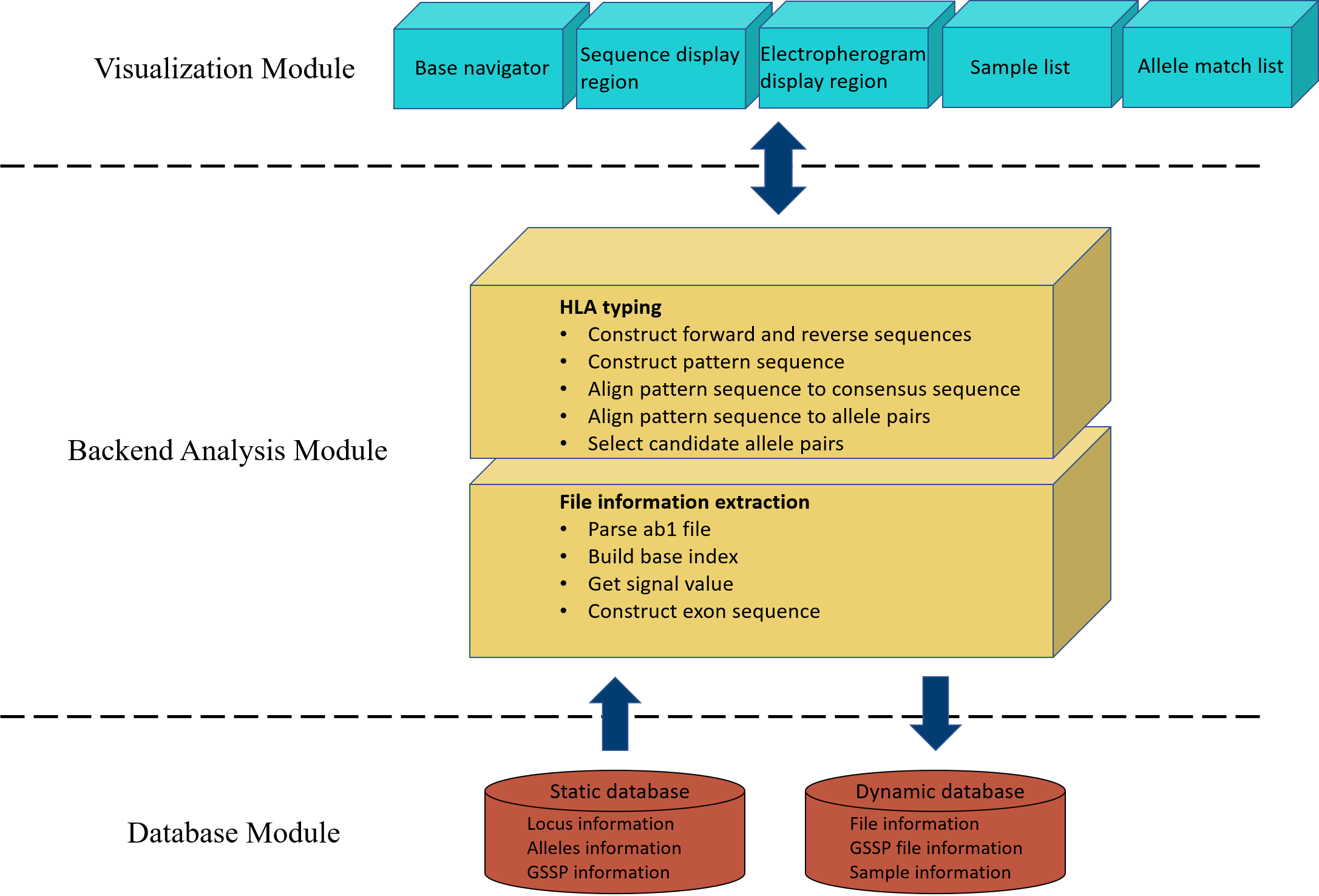

Figure S1. Overview of SOAPTyping architecture. The static database is the basic support for the backend analysis module. The dynamic database stores intermediate data generated by the backend analysis module so that users could get back to states of former analysis even after they have shutdown SOAPTyping. The backend analysis module aims to produce data for visualization module and provide candidate allele pairs for selection. The visualization module is designed for exhibition and interactive operation. It also interacts with the backend analysis module. For example, SOAPTyping will repeat analysis of a specific sample when users edit sites on the interface.

#### 1.1 Visualization module

The visualization module is designed to create interactive interface, including multiple panes of SOAPTyping’s main window. The main window consists of panes of Sample List, Base Navigator, Allele Match List, Sequence Display Region, Electropherogram Display Region and Toolbar (Figure S2).

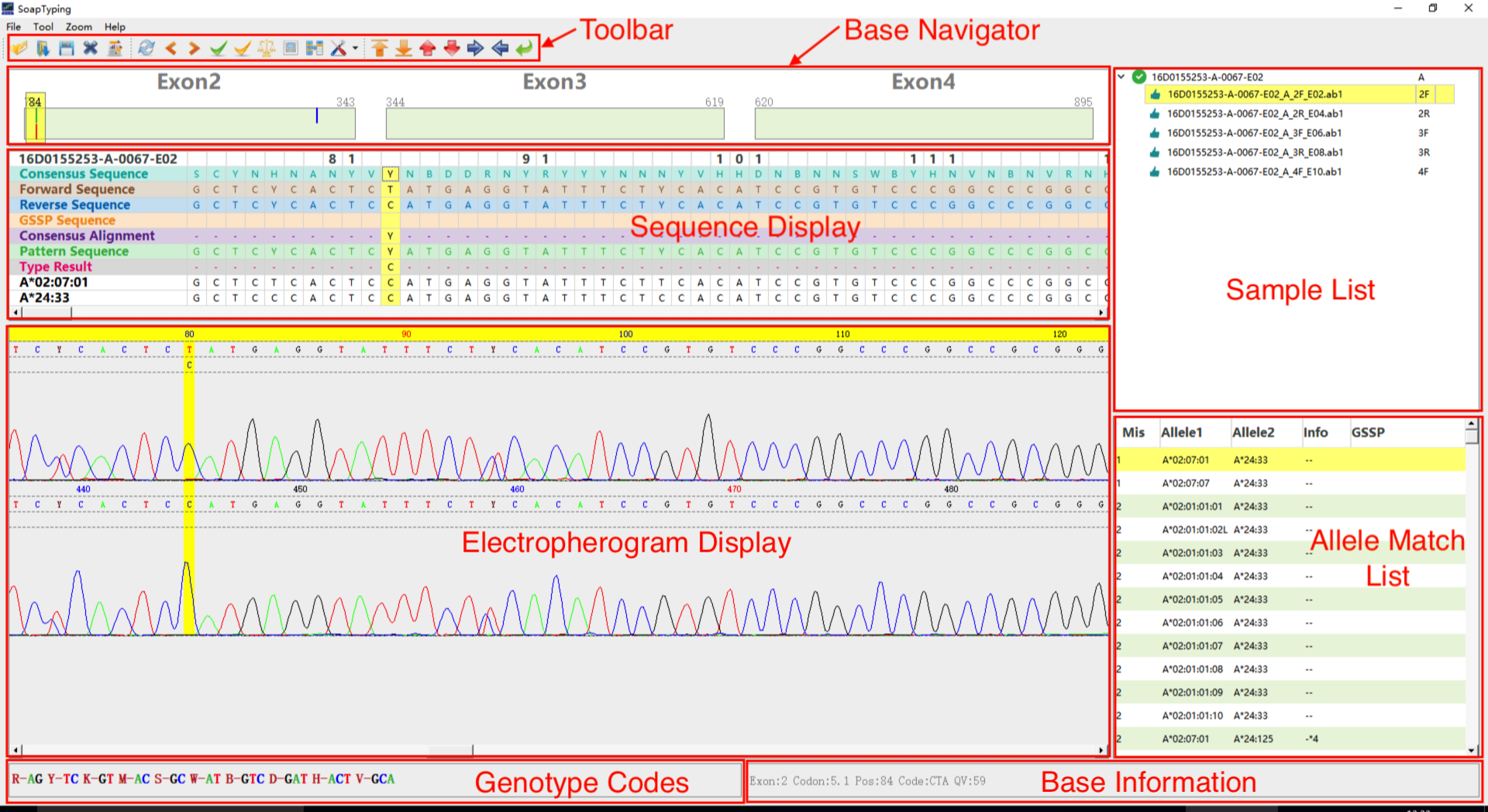

Figure S2. The main window of SOAPTyping.

1.1.1 The pane of Sample List organizes the input files into a tree structure based on the sample name, in which nomenclature is given in the following Section 2.1. Involved icons and their detailed explanations are shown in Table S2.

Table S2. Icons involved in the pane of Sample List

| Icons | Descriptions |
| --- | --- |
| 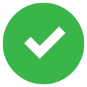 | No mismatches exist between pattern sequence and the allele pair. The allele pair is a common type. |
| 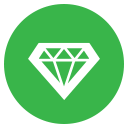 | No mismatches exist between pattern sequence and the allele pair. The allele pair is a rare type. |
| 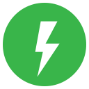 | No mismatches exist between pattern sequence and the allele pair. The quality of the sequence file is poor. |
| 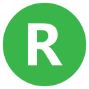 | No mismatches exist between pattern sequence and the allele pair. The sequence file is marked as reviewed. |
| 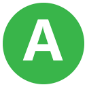 | No mismatches exist between pattern sequence and the allele pair. The sequence file is marked as approved. |
| 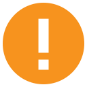 | Mismatches exist between pattern sequence and the allele pair. |
| 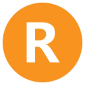 | Mismatches exist between pattern sequence and the allele pair. The sequence file is marked as reviewed. |
| 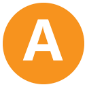 | Mismatches exist between pattern sequence and the allele pair. The sequence file is marked as approved. |
| 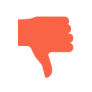 | The quality of the sequence file is poor. |
| 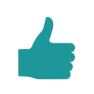 | The quality of the sequence file is satisfactory. |
| 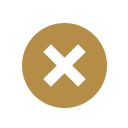 | The sequence file cannot be analyzed. |
| 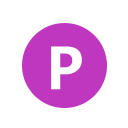 | The sequence file is marked as pending. |

1.1.2 The pane of Base Navigator highlights mismatched positions so that users can skip to such positions quickly by clicking on the color bar, in which detailed descriptions are presented in Table S3. The pane of Base Navigator could be divided into two parts. The upper area displays the difference between the forward and backward sequence as well as the discrepancies between the pattern and consensus sequence. In the meantime, it also tracks the edited or filtered position. The lower area shows mismatches between the pattern sequence and its best matched allele, which are only displayed when a specific allele pair is chosen.

Table S3. Colors and their meanings in the pane of Base Navigator

| Color | Descriptions |
| --- | --- |
| Blue | The forward and backward sequence are compatible but different. |
| Red | The pattern sequence is not compatible with consensus sequence and there exist mismatches between pattern sequence and the chosen allele pair. |
| Green | The forward and backward sequence are not compatible. |
| Gray | The base has been edited. |

1.1.3 The pane of Allele Match List displays possible typing results sorted following the order of number of mismatched sites. Detailed explanations of designed columns are listed in Table S4.

Table S4. Detailed columns showed in the pane of Allele Match List

| Columns | Descriptions |
| --- | --- |
| Mis | Number of mismatches between the allele pair and the pattern sequence. Note that ‘*’ indicates indel(s) exist(s) among at least one of the typed alleles. Such candidate allele pairs could be removed by a double-click. |
| Allele1 | The first candidate allele |
| Allele2 | The second candidate allele |
| Info | Information of allele pairs. A single “r” shows one of the candidate alleles is a rare type and thus double “r” means both candidate alleles are rare types. Symbol of “-” indicates no missing exons used to analyze in candidate alleles. Symbol of “*” followed with a number indicates a specific exon region is missing in the candidate allele. |
| GSSP | “Yes” means GSSP information is available |

1.1.4 The pane of Sequence Display (Table S5), from top to bottom, is comprised of several tracks including ‘Sample and Position’, ‘Consensus Sequence’, ‘Forward Sequence’, ‘Reverse Sequence’, ‘GSSP Sequence’, ‘Consensus Alignment’, ‘Pattern Sequence’, ‘Type Result’ and sequences of the allele pair.

Table S5. Descriptions of each rows in the pane of Sequence Display

| Rows | Descriptions |
| --- | --- |
| 1 | Sample name listed at the beginnings, followed by position information |
| 2 | Consensus Sequence showing the consensus sequence of the locus being analyzed. |
| 3 | Forward sequences that are parsed from the input electropherogram files. |
| 4 | Reverse sequences that are parsed from the input electropherogram files. |
| 5 | GSSP sequences that are parsed from the input electropherogram files. |
| 6 | Consensus Alignment showing alignment results of the pattern sequence and the consensus sequence. |
| 7 | Pattern Sequence showing the combination of the forward and reverse sequence. For example, if a site in the forward sequence is ‘A’, the corresponding site in the reverse sequence is ‘R’, which means this site is a heterozygous site of ‘A’ and ‘G’, in which case the combination of this site is ‘R’. |
| 8 | Type results showing different alleles compared to the chosen allele pair in the pane of Allele Match List. |
| 9 | Sequence of the first allele of the chosen candidate allele pair in the pane of Allele Match List |
| 10 | Sequence of the second allele of the chosen candidate allele pair in the pane of Allele Match List |

1.1.5 The pane of Electropherogram Display Region (Figure S2) displays the electropherogram of the forward sequence, the reverse sequence and the GSSP sequence, so that users can edit bases in this region.
1.1.6 The pane of Toolbar integrates some useful functions and information. The detailed function of each icon is listed in Table S6. Corresponding degenerate bases as well as signal information of the chosen base are shown at the bottom of the UI.

Table S6. Descriptions of icons in the pane of Toolbar

| Icons | Descriptions |
| --- | --- |
| 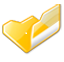 | Open new files |
| 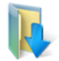 | Load saved files |
| 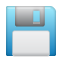 | Save |
| 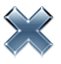 | Delete file in the pane of Sample List |
| 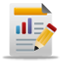 | Export the final report |
| 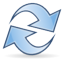 | Reset the file as initial status |
| 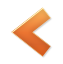 | Jump to the previous mismatched locus |
| 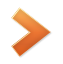 | Jump to the next mismatched locus |
| 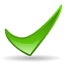 | Set all files status as “Approved” |
| 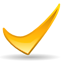 | Set all files status as “Reviewed” |
| 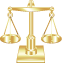 | Allele pair alignments |
| 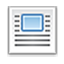 | Multiple alleles alignment |
| 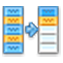 | Update databases |
| 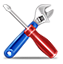 | Set parameters of the software, such as number of threads, exon region needed to be trimmed |
| 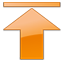 | Y range zoom of the electropherogram (+) |
| 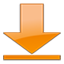 | Y range zoom of the electropherogram (-) |
| 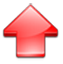 | Heighten the electropherogram |
| 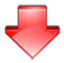 | Shorten the electropherogram |
| 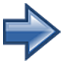 | Widen the electropherogram |
| 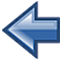 | Narrow the electropherogram |
| 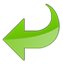 | Reset the electropherogram |

#### 1.2 Database module

The database module mainly includes two kinds of databases, which are static and dynamic. The static database contains information that is not changed during runtime of SOAPTyping, which is composed of tables designed to store details related to HLA loci, alleles, exons, genes and GSSPs. The table of loci stores consensus sequence and available exons that can be analyzed for each locus. The table of alleles stores HLA-I and HLA-II allele sequences and particular details of each allele, including rare variants and indels. The table of GSSP information table stores the GSSPs made for wanted exon regions, inserted sites and inserted bases. These tables could be manually updated (see below at Section 2.9 for database updates). The dynamic database also applies different tables to store intermediate data generated during runtime, including tables of files, GSSP and samples. The dynamic tables of files and GSSPs store information that are required to construct sequence electropherogram. The table of sample mainly stores sequences parsed from input files and typing results of each sample.

***1.2.1	Static database***

Table S7. alleleTable

| **alleleName** | Store the allele name (eg: A*01:01:01:01) |
| --- | --- |
| **alleleSequence** | Store the allele sequence |
| **geneName** | Store the gene name |
| **isRare** | Store the result of rare type (0: normal 1:rare) |
| **isIndel** | Store the result of indel (0:normal 1:indel) |
| **indelPosition** | Store the indel position |
| **indelInfo** | Store the information of indel(eg: Delete 'N') |

Table S8. gsspTable

| **gsspKey** | Store gssp key which consist of gsspName, _,geneName |
| --- | --- |
| **gsspName** | Store gssp name |
| **geneName** | Store gene name |
| **exonIndex** | Store exon index |
| **rOrF** | Store the direction |
| **position** | Store the gssp position |
| **base** | Store the base of gssp position |

Table S9. geneTable

| **geneName** | Store gene name |
| --- | --- |
| **geneSequence** | Store gene sequence |
| **exonCount** | Store the exon count |
| **exonPositionIndex** | Store the exon pos array(eg: 0:73:343:619:895:1012:1045:1093:1098) |
| **geneClasses** | Store the gene class(eg: 0:192:194:224:226:240:242) |
| **availableExon** | Store the exon regions (eg: 123456) |

***1.2.2	Dynamic database***

Table S10. fileTable

| **fileName** | Store the name about the input ab1 file |
| --- | --- |
| **sampleName** | Store the sample name about the input ab1 file |
| **geneName** | Store the gene name about the input ab1 file |
| **exonIndex** | Store the exon index about the input ab1 file |
| **usefulSequence** | Store the alignment sequence about the input ab1 file |
| **baseSequence** | Store the original sequence about the input ab1 file |
| **basePosition** | Store the original position about the input ab1 file |
| **baseQuality** | Store the original quality about the input ab1 file |
| **baseASignal** | Store the original A signal about the input ab1 file |
| **baseTSignal** | Store the original T signal about the input ab1 file |
| **baseGSignal** | Store the original G signal about the input ab1 file |
| **baseCSignal** | Store the original C signal about the input ab1 file |

Table S11. gsspFileTable

| **fileName** | Store the name about the input gssp file |
| --- | --- |
| **sampleName** | Store the sample name about the input gssp file |
| **geneName** | Store the gene name about the input gssp file |
| **exonIndex** | Store the exon index about the input gssp file |
| **usefulSequence** | Store the alignment sequence about the input gssp file |
| **baseSequence** | Store the original sequence about the input gssp file |
| **basePostion** | Store the original position about the input gssp file |
| **baseQuality** | Store the original quality about the input gssp file |
| **baseASignal** | Store the original A signal about the input gssp file |
| **baseTSignal** | Store the original T signal about the input gssp file |
| **baseGSignal** | Store the original G signal about the input gssp file |
| **baseCSignal** | Store the original C signal about the input gssp file |

Table S12. sampleTable

| **sampleName** | Store the name about sample |
| --- | --- |
| **geneName** | Store the gene name about sample |
| **consensusSequence** | Store the consensus sequence which form genetable |
| **forwardSequence** | Store the forward sequence which form filetable |
| **reverseSequence** | Store the reverse sequence which form filetable |
| **patternSequence** | Store the pattern sequence which merge forward sequence and reverse sequence |

#### 1.3 Backend analysis module

The backend analysis module includes two submodules, which are used to perform file information extraction and HLA typing. The utility of the extraction module is to parse input electropherogram files to obtain base sequences. The HLA typing module aims to generate candidate allele pairs. Specifically, it first constructs forward and reverse sequences by assembling different exon sequences that are acquired orderly with utilization of the extraction module. Then the forward and reverse sequences are combined to form a pattern sequence, which is next aligned to the consensus sequence of loci and alleles in the static database. Finally, all candidate allele pairs are collected and sorted according to the counts of the mismatched sites they harbor.

***1.3.1	Base calling***

After ABIF files are loaded to extract needed information, SOAPTyping obtains the details of base sequence, maximum signal position, quality value and base signal values for each A/T/C/G base. To achieve identification of heterozygotes and homozygotes, a peak range of each base is calculated using following formula. The $R_{low}$ and $R_{high}$ is the low and high range of the current base, ${position}_{i}$ is the signal position of the current base peak, while ${position}_{i-1}$ and ${position}_{i+1}$ are the signal position of the previous and next base peak.

$$R_{low}={position}_{i}-\frac{{position}_{i}-{position}_{i-1}}{2},(1)$$

$$R_{high}={position}_{i}+\frac{{position}_{i+1}-{position}_{i}}{2},(2)$$

Next, SOAPTyping will search to find if there exists another peak during this range. Is another peak exists with a signal value greater than 0.3 times the maximum signal within 4 units of distance, such a position will be determined as heterozygous genotypes. Homozygotes will be determined if only one peak exist during this range. The inferred genotypes are presented following the code standard of IUPAC-IUB.

***1.3.2	Sequence alignment and HLA typing***

As variants in the exon regions are considered, SOAPTyping aligns the sanger sequencing sequence to the exon sequence from the IMGT/HLA database using a modified semi-global alignment method. The semi-global alignment method does not penalize gaps at the beginning and/or end of an alignment. The main adjustment of our semi-global alignment method is that comparisons of one degenerated base will be considered as comparisons of two independent alleles derived from that degenerated base, which is shown in the Formula 3. For example, comparisons between degenerated bases of A, R and G and reference A will end up with scores of 2, 1 and 0, respectively.

$$\mathrm{Score}\left( {seq1}_{i}, {seq2}_{j} \right)=\left\{ \begin{aligned} 2, when 2 alleles match \\ 1, when 1 allele match \\ 0, mismatch \\ -1, indel \end{aligned}, (3) \right.$$

Afterwards, SOAPTyping will merge alignment results based on multiple input files. In the merging process, the differences of forward and reverse sequence and differences of the sample sequence and IMGT/HLA types are stored in the dynamic database. Users have access to recorded difference at the main UI. Meanwhile, users can edit mismatched bases at the pane of Electropherogram Display Region, following by SOAPTyping’ automated analysis repeatedly.

### 2. Best practices / propositional workflow

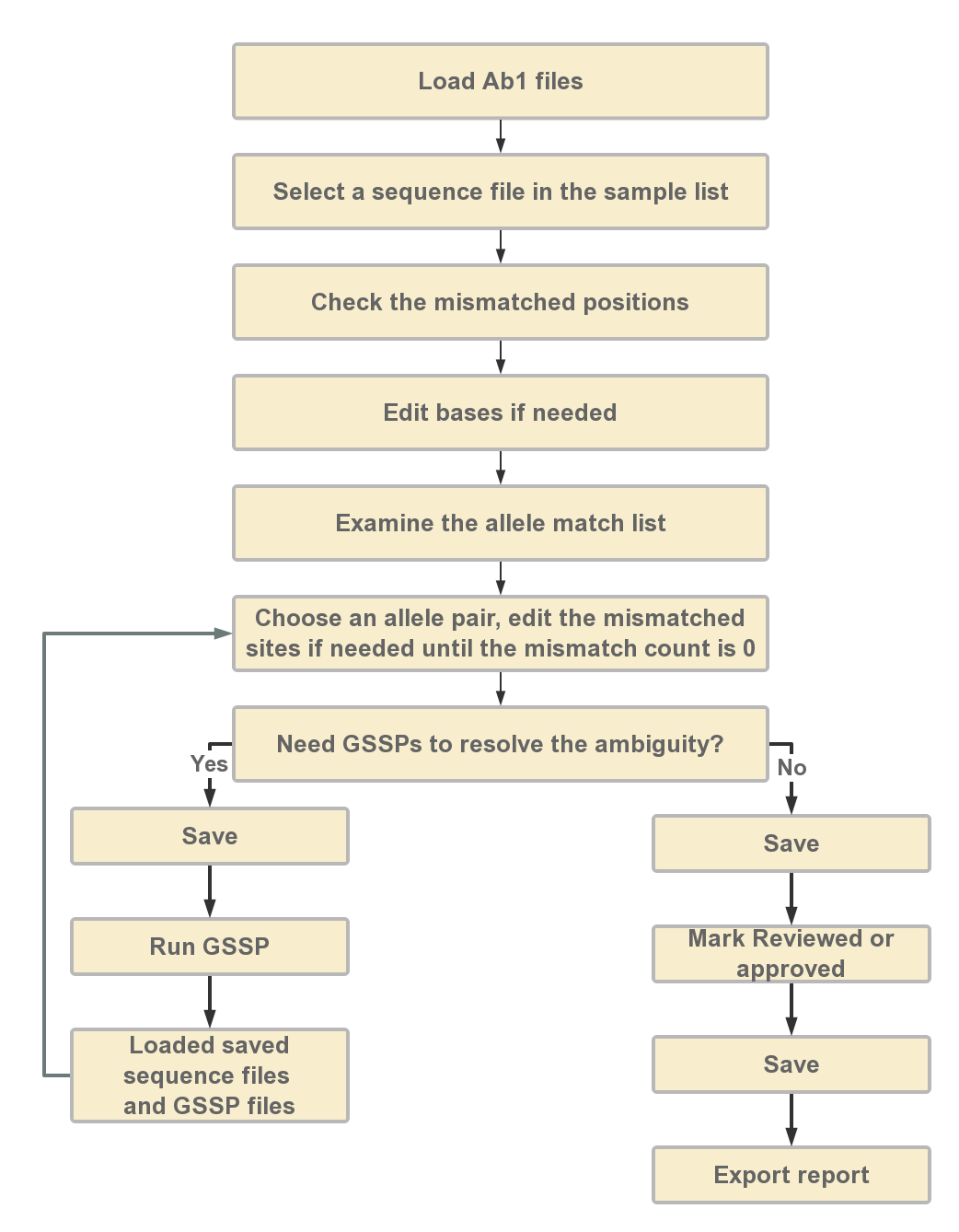
The best practices and propositional workflow are demonstrated in Figure S3. The details of each step are described below.

Figure S3. Best practices and proposed workflow for SOAPTyping.

#### 2.1 Load ABIF files

Users can click the “open new file” icon at toolbar to load the input files. During the loading process, information of imported files is shown in the ‘Open File’ window (Figure S4). To utilize the automatic extraction function, the file name should formatted as: [sample name]_[gene]_[exon region and strand direction]_[sequencer ID] (e.g., RB16002353_A_2R_A02 for sample name: RB16002353, gene: A, exon: 2, strand direction: Reverse, sequencer id: A02). For GSSP file, the file name should formatted as: [sample name]_[gene]_[GSSP name]_[sequencer ID]. If ambiguous nominations of input files cause improper information extraction, users can modify names of input files by choosing correct information from the corresponding drop-down list. User can also delete the files by choosing the file in front of file details and click “Delete Chosen” button. After reassuring files information, users can performance the analysis by click the “Analysis” button.

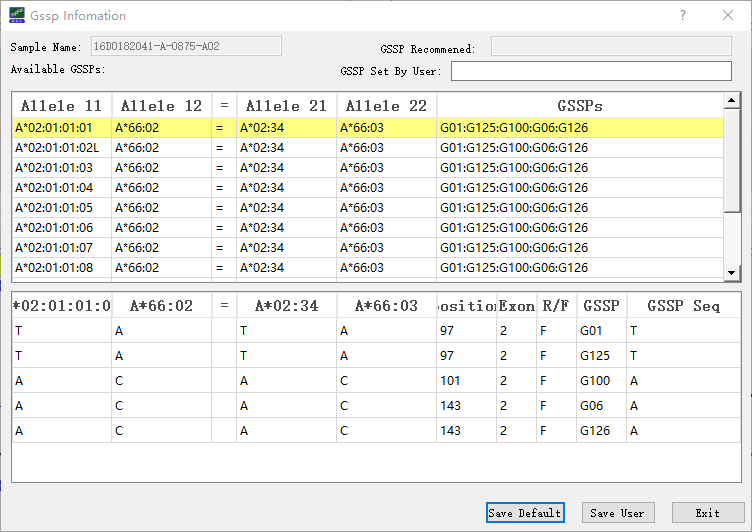

Figure S4. Loading input file.

#### 2.2 Select sequence files in Sample List

Afterwards, imported sequence files are integrated based on sample name and gene, and displayed in the pane of Sample List. If a sample is selected, sequence information, electropherogram and candidate allele pairs of this sample will be displayed in the corresponding region of the main window (Figure S2).

#### 2.3 Check mismatched positions

Two kinds of mismatch, compatible and non-compatible mismatches, are considered in SOAPTyping. For example, ‘R’ (heterozygotes of ‘A’ and ‘G’) and ‘A’ are compatible mismatch and ‘A’ and ‘G’ are a non-compatible mismatch. Users need first to disambiguate the non-compatible sites in forward and reverse sequence, and then disambiguate the non-compatible sites in pattern sequence and consensus sequence. These mismatched sites are pointed out in upper area of the pane of Base Navigator. Users can skip to those positions by clicking on the indication bar and edit bases if needed after examining the electropherogram. To edit a base, click on the site on the electropherogram and input the correct base. Once a base is edited, SOAPTyping will reanalyze this sample and update the allele match list and skip to next mismatched position automatically. If users would like to perform analysis after all mismatched bases are edited, choose ‘Edit Multi’ from the ‘Zoom’ list.

#### 2.4 Examine the allele match list

By selecting a suitable allele pair in the pane of Allele Match List, the corresponding sequences of this allele pair will be shown in the pane of Sequence Display Region. Mismatched sites between pattern sequence and allele pair sequences are highlighted in red in ‘Type Result’ row and shown in the lower area in the pane of Base Navigator. Usually, there are one or serval allele pairs matched to the pattern sequence without any mismatches. But ~0.5% samples may not have a perfect match between the pattern sequence and allele pair sequences in the HLA databases due to the new types or other reasons. Users should make sure the combination of allele pair sequences are perfect matched to pattern sequence before generating results.

#### 2.5 Save and import unfinished jobs

SOAPTyping offers two solutions to save results. One solution is to save results of an individual sample by a right-click on this sample in the pane of Sample List and then choosing options from “Quick Save”, “Quick Save and Clear” or “Quick Save by Date”. Another solution is clicking on the “Save” button in the toolbar to save analysis outcome of multiple samples. During the saving process, a “Result” directory will be created in the work directory at the first time. Under the “Result” directory, each sample is assigned a subdirectory to save the original sequence files and intermediate data that are produced during analysis. For subsequent analysis, users can import saved files by clicking on the “Load” button (Figure S3). SOAPTyping will then search all analysis files of the day and import them, in which the search scope also can be determined by date and names of ABIF files.

#### 2.6 GSSP prediction system

GSSP prediction system is used to resolve ambiguity, in which ambiguity means there are multiple candidate allele pairs that have no mismatches. To use a new GSSP, users need to add it to the database (see section 2.9 for database updates), otherwise SOAPTyping cannot recognize it. Suitable GSSPs are found in SOAPTyping if an allele pair is marked as ‘Yes’ in the ‘GSSP’ column in the pane of Allele Match List. To check GSSP information, users could right-click on the allele pair and choose ‘Show GSSP Z Code’. After resequencing to obtain ABIF files of GSSPs, users could import corresponding sequence files applied to resolve the ambiguity. SOAPTyping will search the candidate allele pairs for each GSSP file and users need to choose a suitable allele pair and edit bases if mismatched sites exist. Finally, users could click on ‘Combined’ under the sample will show the ultimate HLA typing result after all GSSP files have been examined.

#### 2.7 Mark the sample analysis stat

Once sequence files of a sample are imported, SOAPTyping will automatically mark samples status based on sequencing quality of input files and match conditions of pattern sequence and candidate allele pairs. An icon indicating current status is shown before the sample name in the pane of Sample List. Users can also mark the sample as ‘Pending’, ‘Reviewed’ and ‘Approved’ in accordance with the analysis progress. When a defective sample comes up or analysis is not complete, it may be suitable to mark the sample as ‘Pending’ for subsequent analysis. If the typing result is reviewed, such a sample can be marked as ‘Reviewed’. If typing results is affirmed, such a sample can be labeled as ‘Approved’. The status also can be canceled by choosing ‘Unlock’. All samples can be marked as ‘Reviewed’ or ‘Approved’ simultaneously by clicking on the corresponding button in the toolbar while marking ‘Pending’ is only allowed for single sample.

#### 2.8 Export reports

To export reports, users could click on the corresponding button in the pane of Toolbar and the window of ‘Produce report’ will appear. Users can name reports and the output directory. When ‘Ignore Indel’ is selected, alleles that contain indels would be excluded. The ‘Allele Count’ option is to control numbers of the output alleles, otherwise all alleles without mismatches will be exported by default. A report example is demonstrated in Figure S5.

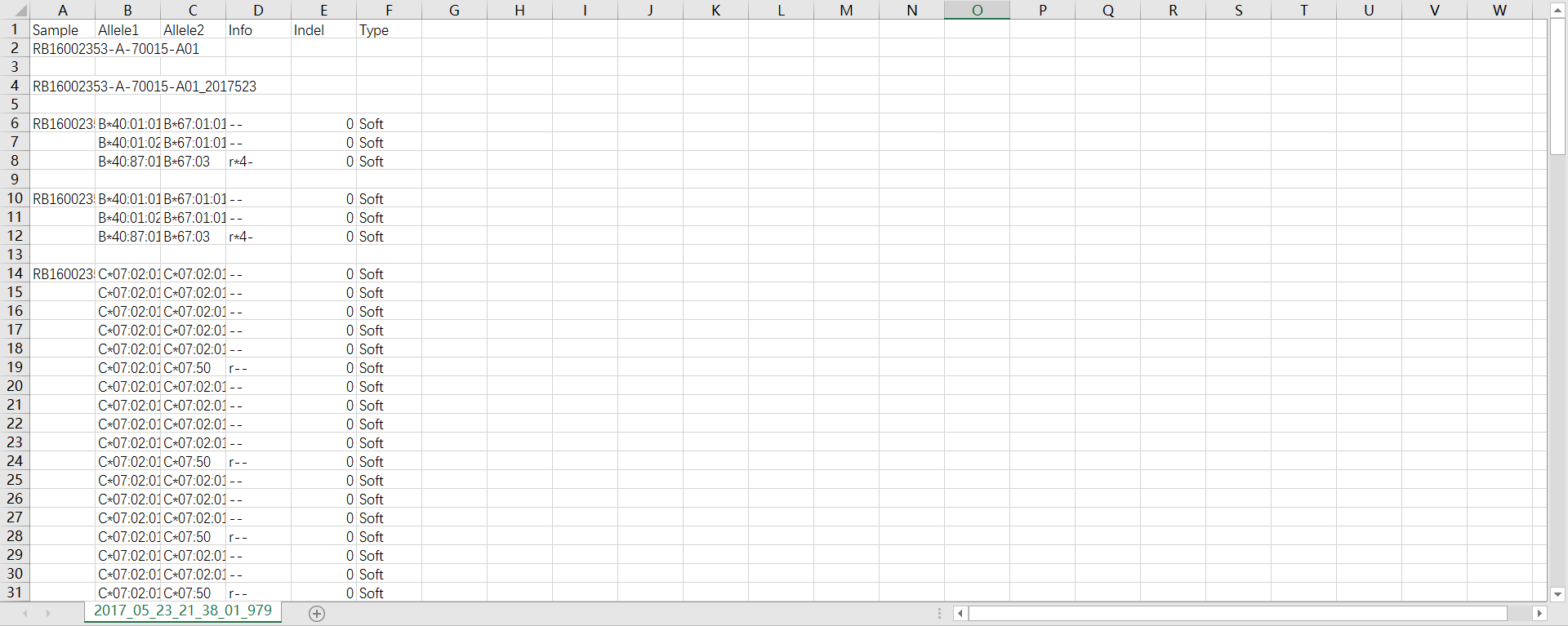

Figure S5. An example exported report. In the ‘Type’ column, ‘Soft’ indicates no GSSP is used and ‘Filter’ means the opposite.

#### 2.9 Database update

SOAPTyping offers database update function to cater to the situation of frequent update of HLA alleles. Before updating the database, users could download the latest release of compressed alignments file (‘zip’ format, e.g., Alignments_Rel_3260.zip) of IMGT/HLA database via its FTP directory (ftp://ftp.ebi.ac.uk/pub/databases/ipd/imgt/hla/). After decompressing the downloaded zip files, users can get an ‘alignments’ folder which contains genomics sequence alignments, nucleotide sequence alignments and protein sequence alignments files for each locus, then select the nucleotide sequence alignments files and perform sequence format conversion using the scripts we provided on our website. This step generates several text files, including ‘geneTable’, ‘alleleTable’ and ‘labAlignTable’. These files correspond ‘Gene File’, ‘Allele File’ and ‘Lab Align File’ respectively as required in the database update window (Figure S6). Details of these update files are illuminated at Table S13 and Figure S7.

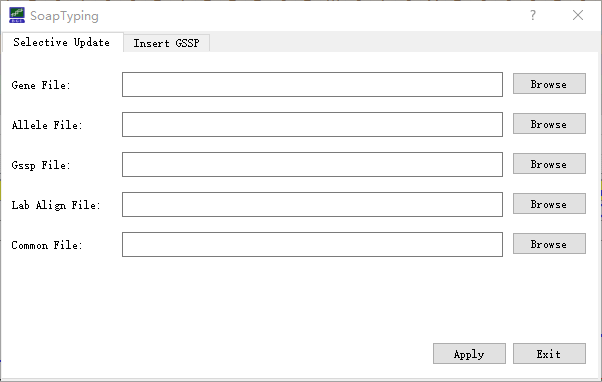

Figure S6. Files required for database updates

Table S13. Details of files required for database update

| Filename | Content |
| --- | --- |
| geneTable.txt | New HLA molecules consensus sequences |
| oldGeneTable.txt | HLA molecules consensus sequences that needed to be update |
| alleleTable.txt | HLA allele sequences |
| gsspTable.txt | GSSP information |
| labAlignTable.txt | Alleles alignment information |
| exonTrimTable.txt | Exon regions excluded from analysis |

a)

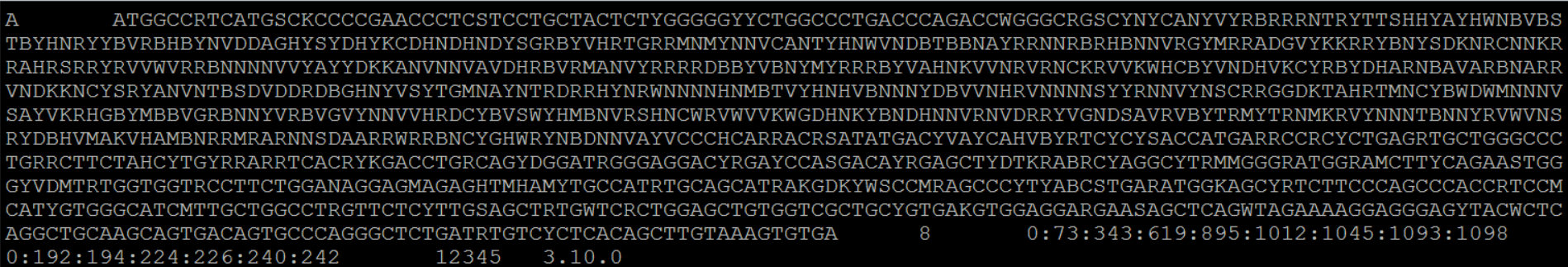

b)

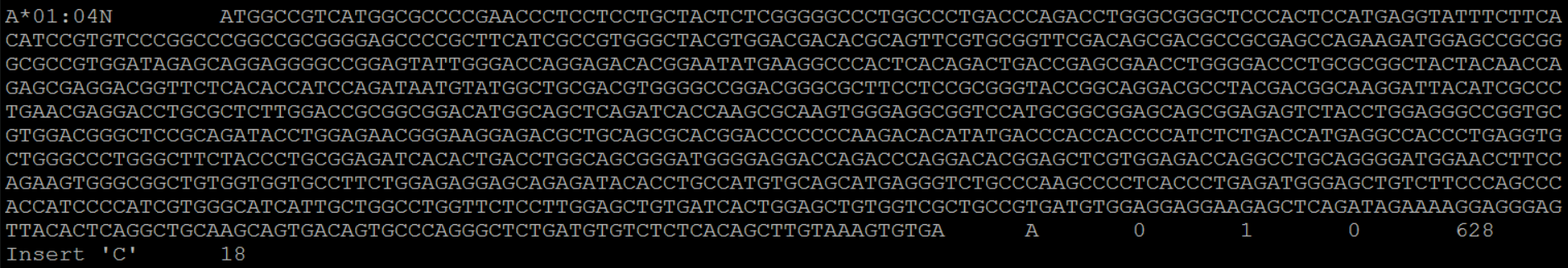

c)

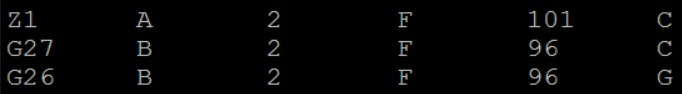

d)

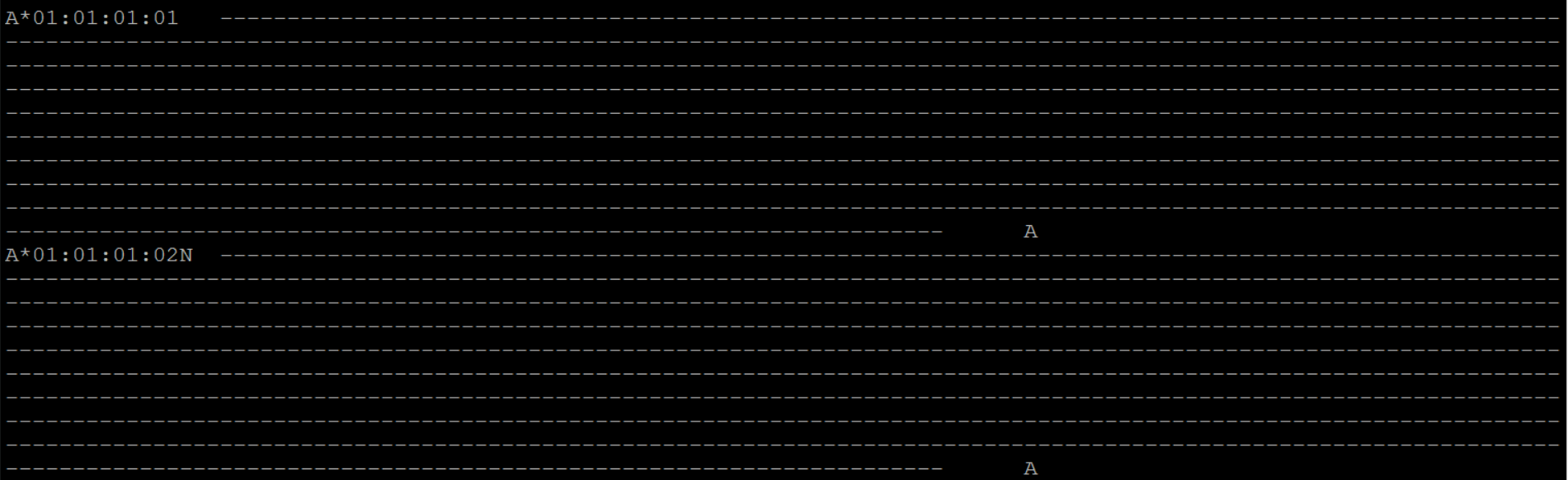

e)

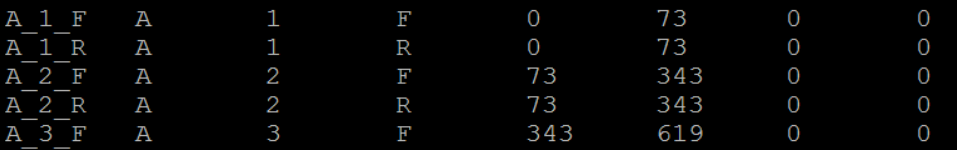

Figure S7. Screenshots of example files needed for database update.

a) geneTable and oldGeneTable. Columns from left to right show HLA genes, consensus sequence, number of exons, exon intervals, gene classes, available exons for analysis and the version, which concurs with the IGMT
b) alleleTable. Columns from left to right show allele names, allele sequence, loci, if it is rare (0 means no otherwise yes), if it contains indels, gene classes, indel positions, indel descriptions, exon regions that has no ‘*’.
c) gsspTable. Columns from left to right show GSSP names, loci, exon index, forward or reverse strand, positions, bases.
d) labAlignTable. Columns from left to right show allele names, alignment status and loci. Specifically, each locus have a standard sequence and all alleles in the same locus are aligned to this sequence. For example, A*01:01:01:01 is the standard sequence for locus A. Symbol of “-“ means the base is correctly matched to the standard sequence and the aligned base will be shown up if not matched and ‘*’ means a base is missed.
e) exonTrimTable. Columns from left to right show loci, exon index, forward strand or reverse strand, start positions, end positions, bases trimmed of the left side and bases trimmed of the right side of the exon.

Additionally, if users want to update the GSSP information, text file contains GSSP information should be prepared manually, in which examples are illustrated in Figure S8. Another way to update GSSPs is to fill the GSSP information in the corresponding fields in the window of ‘Insert GSSP’ (Figure S9).

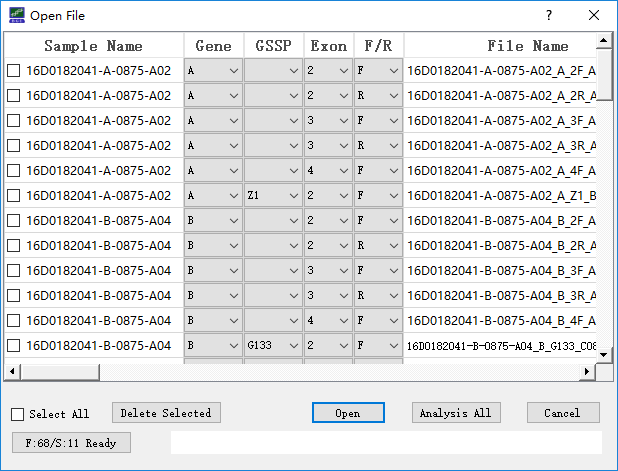

Figure S8. The GSSP column show the GSSPs that can discern the ambiguity and the detailed information is shown below.

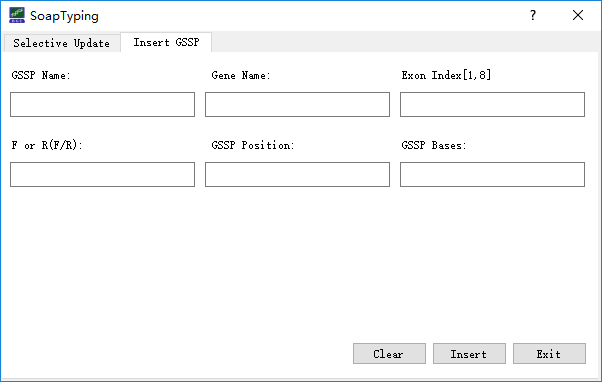

Figure S9. Files required for GSSP database update

#### 2.10 Utilities

SOAPTyping supplies a set of utilities to assist HLA typing, which are listed as followings. (1) Setting analysis threads. Click on the ‘Setting’ button (Figure S2) and choose ‘Set Thread’ to set a suitable number of threads to accelerate analysis. (2) Customizing the exon region for analysis. Click on the ‘Setting’ button and choose ‘Set Exon Trim’ to exclude bases of both ends of the exon, in which excluded parts are not considered in analysis. (3) Alleles alignment (Figure S10a). Click on the ‘Allele Alignment’ button to invoke the alignment tool and choose a locus from the ‘Genes’ drop-down list. All available alleles of this locus will be listed in the panel below (Figure S10b). The alignment result will be displayed after alleles of interest are chosen.

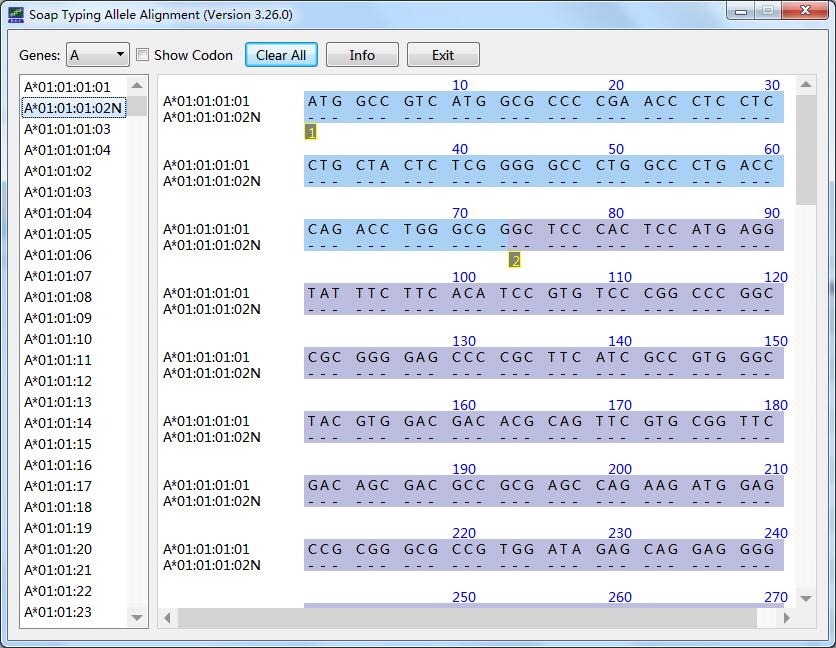

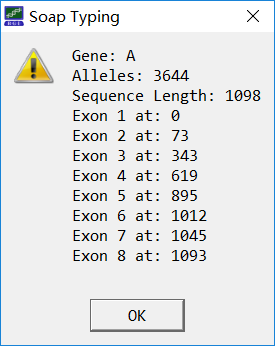

a) b)

Figure S10. Allele alignment tool. a) Alleles of a locus will be aligned to a standard sequence. ‘-’ means a base is correctly matched, otherwise, mismatched bases are shown. ‘*’ indicates a base is missing. The alignment result is relevant to the database file ‘labAlignTable’ (Table S6, Figure S7); b) Clicking ‘Info’ will display the statistic information of the locus.

### 3. Verification using test data

### 3.1 Test data

Our test data contains 36 samples initiated for external quality assessments with the University of California Los Angeles (UCLA) International HLA DNA Exchange (Los Angeles, CA, USA). Genomic DNAs with known HLA typing results were obtained from UCLA and amplified using locus-specific primers. All samples have been typed for HLA-A, -B, -C, -DRB1, and -DQB1 by Sanger sequencing using a 3730XL DNA Analyzer (Applied Biosystems, Foster City, CA). Sequencing reaction was performed using the BigDye® Terminator v3.1 Cycle Sequencing Ready Reaction Kit (Applied Biosystems). Sequencing reaction was performed using the BigDye® Terminator v3.1 Cycle Sequencing Ready Reaction Kit (Applied Biosystems). The Sanger sequencing strategy involved amplification spanning exons 2, 3 and 4 for HLA-A, -B, -C, and two separate amplicons for exons 2 and 3 for HLA-DQB1 and -DPB1 (Table S1). Group specific sequencing primers (GSSP) were applied to resolve ambiguities. The test data have been deposited in the CNSA (https://db.cngb.org/cnsa/) of CNGBdb with an accession code CNP0000512.

### 3.2 Results

The sequence was analyzed with SOAPTyping. The typing results were compared to the consensus based on high resolution provided by UCLA. The consistency of SOAPTyping in typing HLA alleles at four-digit was verified to be accurate at the level of 100% (36/36) for HLA-A, -B, -C, -DR and -DQ. The detailed results of 36 tested samples were shown in Table S13.

Table S13. SOAPTyping results of 36 samples from UCLA International DNA Exchange

| SampleNameUcla | Population | A1 | A2 | B1 | B2 | C1 | C2 | DR1 | DR2 | DQ1 | DQ2 |
| --- | --- | --- | --- | --- | --- | --- | --- | --- | --- | --- | --- |
| #867 | BLACK | A*02:01 | A*66:02 | B*44:02 | B*57:03 | C*05:01 | C*06:02 | DRB1*03:01 | DRB1*04:02 | DQB1*02:01 | DQB1*03:02 |
| #868 | HISPANIC | A*01:01 | A*02:01 | B*52:01 | B*57:01 | C*03:03 | C*07:01 | DRB1*01:01 | DRB1*14:06 | DQB1*05:01 | DQB1*03:01 |
| #869 | ASIAN | A*02:01 | A*31:01 | B*27:05 | B*40:06 | C*02:02 | C*08:01 | DRB1*04:01 | DRB1*09:01 | DQB1*03:02 | DQB1*03:03 |
| #870 | CAUCASIAN | A*11:01 | A*25:01 | B*15:01 | B*44:02 | C*03:03 | C*05:01 | DRB1*04:01 | DRB1*13:01 | DQB1*06:03 | DQB1*03:01 |
| #871 | HISPANIC | A*26:01 | A*29:02 | B*27:05 | B*44:03 | C*02:02 | C*16:01 | DRB1*07:01 | DRB1*08:03 | DQB1*02:02 | DQB1*03:01 |
| #872 | CAUCASIAN | A*03:01 | A*03:01 | B*07:02 | B*44:02 | C*05:01 | C*07:02 | DRB1*04:01 | DRB1*15:01 | DQB1*06:02 | DQB1*03:01 |
| #873 | HISPANIC | A*03:01 | A*03:01 | B*07:02 | B*45:01 | C*06:02 | C*07:02 | DRB1*04:01 | DRB1*16:01 | DQB1*05:02 | DQB1*03:01 |
| #874 | ASIAN | A*02:07 | A*11:01 | B*15:58 | B*46:01 | C*01:02 | C*01:02 | DRB1*14:54 | DRB1*15:01 | DQB1*05:02 | DQB1*05:02 |
| #875 | CAUCASIAN | A*02:01 | A*30:02 | B*15:04 | B*18:01 | C*03:03 | C*05:01 | DRB1*03:01 | DRB1*08:04 | DQB1*02:01 | DQB1*04:02 |
| #876 | CAUCASIAN | A*02:01 | A*66:01 | B*27:05 | B*40:01 | C*01:02 | C*03:04 | DRB1*01:01 | DRB1*04:04 | DQB1*05:01 | DQB1*03:02 |
| #877 | BLACK | A*24:02 | A*34:02 | B*13:02 | B*14:02 | C*06:02 | C*08:02 | DRB1*07:01 | DRB1*14:54 | DQB1*05:03 | DQB1*02:02 |
| #878 | UNKNOWN | A*26:01 | A*26:03 | B*40:02 | B*51:01 | C*03:04 | C*14:02 | DRB1*09:01 | DRB1*14:03 | DQB1*03:01 | DQB1*03:03 |
| #879 | CAUCASIAN | A*02:01 | A*26:01 | B*14:01 | B*55:01 | C*03:03 | C*08:02 | DRB1*07:01 | DRB1*11:03 | DQB1*02:02 | DQB1*03:01 |
| #880 | HISPANIC | A*02:01 | A*26:01 | B*14:02 | B*53:01 | C*01:02 | C*08:02 | DRB1*03:01 | DRB1*13:02 | DQB1*06:09 | DQB1*02:01 |
| #881 | BLACK | A*03:01 | A*33:01 | B*51:01 | B*57:03 | C*01:02 | C*18:02 | DRB1*11:01 | DRB1*11:01 | DQB1*03:01 | DQB1*03:19 |
| #882 | HISPANIC | A*02:01 | A*02:06 | B*07:02 | B*49:01 | C*07:01 | C*07:02 | DRB1*13:02 | DRB1*15:01 | DQB1*06:02 | DQB1*06:04 |
| #883 | ASIAN | A*02:06 | A*24:02 | B*56:01 | B*56:02 | C*01:02 | C*01:02 | DRB1*04:03 | DRB1*09:01 | DQB1*03:02 | DQB1*03:03 |
| #884 | CAUCASIAN | A*29:02 | A*32:01 | B*13:02 | B*44:03 | C*06:02 | C*16:01 | DRB1*07:01 | DRB1*12:01 | DQB1*02:02 | DQB1*03:01 |
| #885 | BLACK | A*25:01 | A*74:01 | B*08:01 | B*15:03 | C*02:10 | C*07:01 | DRB1*03:01 | DRB1*11:01 | DQB1*02:01 | DQB1*03:19 |
| #886 | HISPANIC | A*01:01 | A*25:01 | B*51:01 | B*57:01 | C*06:02 | C*12:03 | DRB1*04:07 | DRB1*13:02 | DQB1*06:04 | DQB1*03:01 |
| #887 | HISPANIC | A*02:01 | A*34:02 | B*40:01 | B*53:01 | C*03:04 | C*06:02 | DRB1*09:01 | DRB1*11:01 | DQB1*02:02 | DQB1*03:01 |
| #888 | ASIAN | A*02:01 | A*02:06 | B*40:01 | B*55:02 | C*04:82 | C*08:01 | DRB1*04:03 | DRB1*08:03 | DQB1*06:01 | DQB1*03:02 |
| #889 | CAUCASIAN | A*01:01 | A*29:02 | B*08:01 | B*45:01 | C*06:02 | C*07:01 | DRB1*04:01 | DRB1*04:01 | DQB1*03:01 | DQB1*03:02 |
| #890 | CAUCASIAN | A*03:01 | A*30:02 | B*14:02 | B*35:01 | C*04:01 | C*08:02 | DRB1*08:06 | DRB1*13:01 | DQB1*06:02 | DQB1*06:03 |
| #891 | BLACK | A*66:02 | A*68:02 | B*39:10 | B*57:03 | C*12:03 | C*18:02 | DRB1*13:02 | DRB1*16:02 | DQB1*05:02 | DQB1*06:09 |
| #892 | HISPANIC | A*02:06 | A*31:01 | B*15:01 | B*39:06 | C*01:02 | C*07:02 | DRB1*08:02 | DRB1*14:06 | DQB1*03:01 | DQB1*04:02 |
| #893 | HISPANIC | A*23:01 | A*30:10 | B*41:01 | B*41:01 | C*06:02 | C*08:02 | DRB1*04:05 | DRB1*11:01 | DQB1*06:02 | DQB1*02:02 |
| #894 | CAUCASIAN | A*01:01 | A*29:02 | B*07:02 | B*07:02 | C*07:02 | C*07:02 | DRB1*15:01 | DRB1*15:01 | DQB1*06:02 | DQB1*06:02 |
| #895 | ASIAN | A*02:01 | A*02:06 | B*40:06 | B*51:01 | C*08:01 | C*15:02 | DRB1*08:02 | DRB1*08:03 | DQB1*06:01 | DQB1*04:02 |
| #896 | CAUCASIAN | A*02:01 | A*03:01 | B*27:02 | B*41:01 | C*02:02 | C*17:01 | DRB1*04:04 | DRB1*13:01 | DQB1*06:03 | DQB1*04:02 |
| #897 | HISPANIC | A*02:05 | A*68:01 | B*48:01 | B*50:01 | C*06:02 | C*08:01 | DRB1*04:04 | DRB1*07:01 | DQB1*02:02 | DQB1*03:02 |
| #898 | BLACK | A*02:01 | A*24:02 | B*14:01 | B*45:01 | C*02:10 | C*16:01 | DRB1*01:02 | DRB1*07:01 | DQB1*05:01 | DQB1*02:02 |
| #899 | UNKNOWN | A*03:01 | A*33:03 | B*18:01 | B*49:01 | C*05:01 | C*07:01 | DRB1*03:02 | DRB1*11:02 | DQB1*03:19 | DQB1*04:02 |
| #900 | CAUCASIAN | A*01:01 | A*02:01 | B*27:05 | B*37:01 | C*02:02 | C*06:02 | DRB1*10:01 | DRB1*14:54 | DQB1*05:01 | DQB1*05:03 |
| #901 | ASIAN | A*03:01 | A*03:01 | B*07:02 | B*35:01 | C*04:01 | C*07:02 | DRB1*01:01 | DRB1*15:01 | DQB1*05:01 | DQB1*06:02 |
| #902 | CAUCASIAN | A*01:01 | A*02:01 | B*08:01 | B*37:01 | C*06:02 | C*07:01 | DRB1*03:01 | DRB1*11:01 | DQB1*02:01 | DQB1*03:01 |
